# A non-canonical role for p27^Kip1^ in restricting proliferation of corneal endothelial cells during development

**DOI:** 10.1101/721621

**Authors:** Dennis M. Defoe, Huiying Rao, David J. Harris, Preston D. Moore, Jan Brocher, Theresa A. Harrison

**Author notes:** New York Eye and Ear Infirmary of Mount Sinai, New York, NY 10003 United States of America. Sarah Cannon Development Innovations, 1100 Dr. Martin L. King Jr. Boulevard, Suite 800, Nashville, TN 37203, United States of America. Competing Interests: The authors have declared that no competing interests exist.

## Abstract

The cell cycle regulator p27^Kip1^ is a critical factor controlling cell number in many lineages. While its anti-proliferative effects are well-established, the extent to which this is a result of its function as a cyclin-dependent kinase (CDK) inhibitor or through other known molecular interactions is not clear. To genetically dissect its role in the developing corneal endothelium, we examined mice harboring two loss-of-function alleles, a null allele (*p27*^−^) that abrogates all protein function and a knockin allele (*p27^CK^*^−^) that targets only its interaction with cyclins and CDKs. Whole-animal mutants, in which all cells are either homozygous knockout or knockin, exhibit identical proliferative increases (∼0.6-fold) compared with wild-type tissues. On the other hand, use of mosaic analysis with double markers (MADM) to produce infrequently-occurring clones of wild-type and mutant cells within the same tissue environment uncovers a roughly three- and six-fold expansion of individual *p27^CK^*^−/*CK*−^ and *p27*^−/−^ cells, respectively. Mosaicism also reveals distinct migration phenotypes, with *p27*^−/−^ cells being highly restricted to their site of production and *p27^CK^*^−/*CK*−^ cells more widely scattered within the endothelium. Using a density-based clustering algorithm to quantify dispersal of MADM-generated clones, a four-fold difference in aggregation is seen between the two types of mutant cells. Overall, our analysis reveals that, in developing mouse corneal endothelium, p27 regulates cell number by acting cell autonomously, both through its interactions with cyclins and CDKs and through a cyclin-CDK-independent mechanism(s). Combined with its parallel influence on cell motility, it constitutes a potent multi-functional effector mechanism with major impact on tissue organization.

## Introduction

The corneal endothelium is a fluid transporting epithelium that is essential for corneal hydration and transparency. Consisting of a monolayer of flat, irregularly-shaped cells with dendrite-like processes, this tissue provides an important physiological interface between the cornea and the adjoining aqueous humor (1-7). A critical determinant of endothelium function is cell density, which in healthy adult humans generally exceeds 2500 cells/mm^2^ (8-9). Because mature corneal endothelial cells (CECs) do not undergo large-scale renewal, cell death due to disease, injury or aging leads inevitably to a decline in cell number (10-14). When cell density falls below approximately 500 cells/mm^2^, a condition referred to as corneal endothelial disease occurs, in which corneal fluid balance is severely compromised and the cornea swells and becomes opaque. Despite the presence under certain circumstances of a corneal wound healing response, in which cell polyploidization and enlargement is triggered (15-16), this and other potential repair mechanisms are apparently insufficient to maintain tissue integrity under severe conditions. In these cases, surgical intervention via transplantation is generally required to correct the resulting endothelial dysfunction (17-19). Thus, in the normal corneal endothelium, tissue structure and function are largely dependent on the continued maintenance of cells generated during the developmental period. For this reason, the mechanisms by which cell number is established and preserved are of fundamental importance.

Developmentally, the endothelium is derived from cranial neural crest cells which, following their migration into the eye, undergo a mesenchymal-epithelial transition to construct an integrated monolayer (20). Studies in rodents have shown that presumptive CECs actively proliferate during the embryological and early postnatal periods but become arrested in the G_1_ phase of the cell cycle shortly thereafter (21). While the mechanisms responsible for limiting cell division are incompletely understood, the cyclin-dependent kinase (CDK) inhibitor p27^Kip1^ (hereafter referred to as p27) is thought to play a central role. In the corneal endothelium of rats and mice, expression of p27 is upregulated during the first two weeks of postnatal life at a time when endothelial cell proliferation declines and monolayer maturation takes place (21-24). Furthermore, in *p27* knockout mice there is both enhanced production of CECs and extension of cell division further into the postnatal period (24).

As a component of the retinoblastoma (Rb) pathway, p27 is a critical modulator of progression through the G_1_ phase of the cell cycle. Initially characterized as a relatively broad-based CDK inhibitor, its most important targets are now recognized to be cyclin E and CDK2 (25). Through simultaneous binding to both proteins, p27 is able to block cyclin-CDK interaction, as well as interfere with ATP binding to the kinase, thus inhibiting catalytic activity (26). Genetically-engineered mice have been particularly informative in outlining the role of this inhibitor in postnatal growth of many tissues (27-31). For example, *p27*-null animals are larger than normal and exhibit multi-organ hyperplasia. Furthermore, the fact that deficient cells are smaller than control cells in some tissues may reflect altered coupling between cell growth and the cell division cycle (27, 32).

In interpreting results from knockout mice, it is often assumed that the sole effect of *p27* gene ablation on proliferation is its interference with operation of the core cell cycle machinery. However, recent evidence has indicated that, in addition to its established role as a cyclin-CDK inhibitor, p27 may also function indirectly as an anti-proliferation factor by restraining mitogenic cell signaling through its interaction with the microtubule-destabilizing protein stathmin (33, 34). Thus, the possibility exists that p27 could be influencing endothelial cell proliferation through both cyclin-CDK-dependent and -independent pathways.

To begin to dissect *p27* gene function in mouse corneal endothelium, we have compared a knockout line (*p27*^−^) with a knockin strain (*p27^CK^*^−^) in which four amino acid substitutions in the *p27* (*cdkn1b*) gene product prevent the interaction of p27 with cyclins and CDK but leave its other functions intact (30, 31, 35). As with previous studies of whole-organism mutants, our results demonstrate very similar proliferative phenotypes for the endothelium in these two types of mice. Importantly, however, use of mosaic analysis with double markers (MADM) to cause sporadic induction of homozygous mutant and wild-type cells in heterozygous monolayers reveals dramatic increases in cell proliferation that are quantitatively different for *p27*-null and *p27^CK^*^−^ lines. While expansion of *p27^CK^*^−^ cells is almost three-fold greater than wild-type cells, that of *p27*-null cells is increased approximately six-fold. These data indicate that p27 exerts its influence on cell division through both cell cycle-dependent and -independent mechanism and that, in both cases, the effect is cell-autonomous. Finally, our evidence showing that the two mutants have opposite effects on cell dispersion within endothelial monolayers confirm previous work documenting the inhibitor’s involvement in cell migration and highlight the fact that p27 is able to regulate proliferation and migration in a coordinate fashion during development.

## Materials and Methods

### Ethics statement

The use of animals was approved by the University Committee on Animal Care (UCAC) at East Tennessee State University (ETSU) and was in compliance with the National Institutes of Health Guidelines for Care and Use of Animals in Research and the ARVO Statement for Use of Animals in Ophthalmic and Vision Research. Mice were housed in the Division of Laboratory Animal Resources at ETSU, a facility accredited by the Association for the Assessment and Accreditation of Laboratory Animal Care (AAALAC), and maintained in an environment of 12-h light/12-h dark.

### Animals

Conventional genetic analysis was carried out using two whole-animal mutant mouse strains, a *p27*-null strain (designated *p27*^−^) with targeted disruption of the entire *p27* coding region and a knock-in line (*p27^CK−^*) in which point mutations abrogate the ability of p27 to bind cyclins and CDKs (27, 30). The null strain (129-*Cdkn1b^tm1Mlf^*/J; Stock #003122) was obtained from The Jackson Laboratory (Bar Harbor, ME). These mice are larger than normal littermates due to extra cell divisions in virtually all lineages. The knockin strain, which has a similar phenotype, was a generous gift of Dr. Michael Dyer (St. Jude Children’s Research Hospital). Hemizygous (*p27^+/−^*) or heterozygous (*p27^+/CK−^*) mice were interbred to generate the *p27*^−/−^ and *p27^CK−/CK−^* animals and wild-type littermates (*p27*^+/+^) used in these experiments. Genotyping was as described previously (27, 30).

For mosaic analysis, the two *p27* mutant strains were used in combination with four other lines: *MADM-6^GR^* (129-*Gt(ROSA)26Sor^tm3(CAG-EGFP/Dsred2)Luo^*/J; Stock #006041) (36), *MADM-6^RG^* (129-*Gt(ROSA)26Sor^tm2(CAG-Dsred2/EGFP)Luo^*/J; Stock #006067) (36), *Hprt-Cre* (37) (129S1/Sv-*Hprt^tm1(CAG-cre)Mnn^*/J Stock #004302) and C57BL/6J (Stock #000664), all from The Jackson Laboratory. *MADM-6^GR^* and *MADM-6^RG^* each carry marker transgenes, targeted to the *Rosa26* locus on chromosome 6, that are reciprocally chimeric for red (R) and green (G) fluorescent proteins (Fig 1). In the case of *MADM-6^GR^* mice, the N terminal coding region of EGFP is combined with the C terminal coding region of DsRed2, while in *MADM-6^RG^* mice the orientation is reversed. To allow enhanced visualization of DsRed2, six copies of the Myc epitope were engineered into the constructs used to produce transgenes so that the expressed protein can be labeled using an anti-c-Myc antibody. Interposed between the N-and C-terminal sequences is an intron within which is embedded a single *loxP* site. Because the intron shifts the reading frame, any proteins produced are nonfunctional. However, when the *MADM-6^GR^* and *MADM-6^RG^* transgenes are present on homologous chromosomes, a Cre-catalyzed interchromosomal recombination event will result in the exchange of C-and N-terminal portions, reconstituting the coding regions for each of the original fluorescent proteins (Fig 1). In these studies, recombinase activity was supplied by a Cre transgene targeted to the X-linked *HPRT* gene, which is expressed ubiquitously. All strains were kept as separate homozygous stocks before MADM analysis and were genotyped by PCR as described (36, 37).

**Figure 1.**
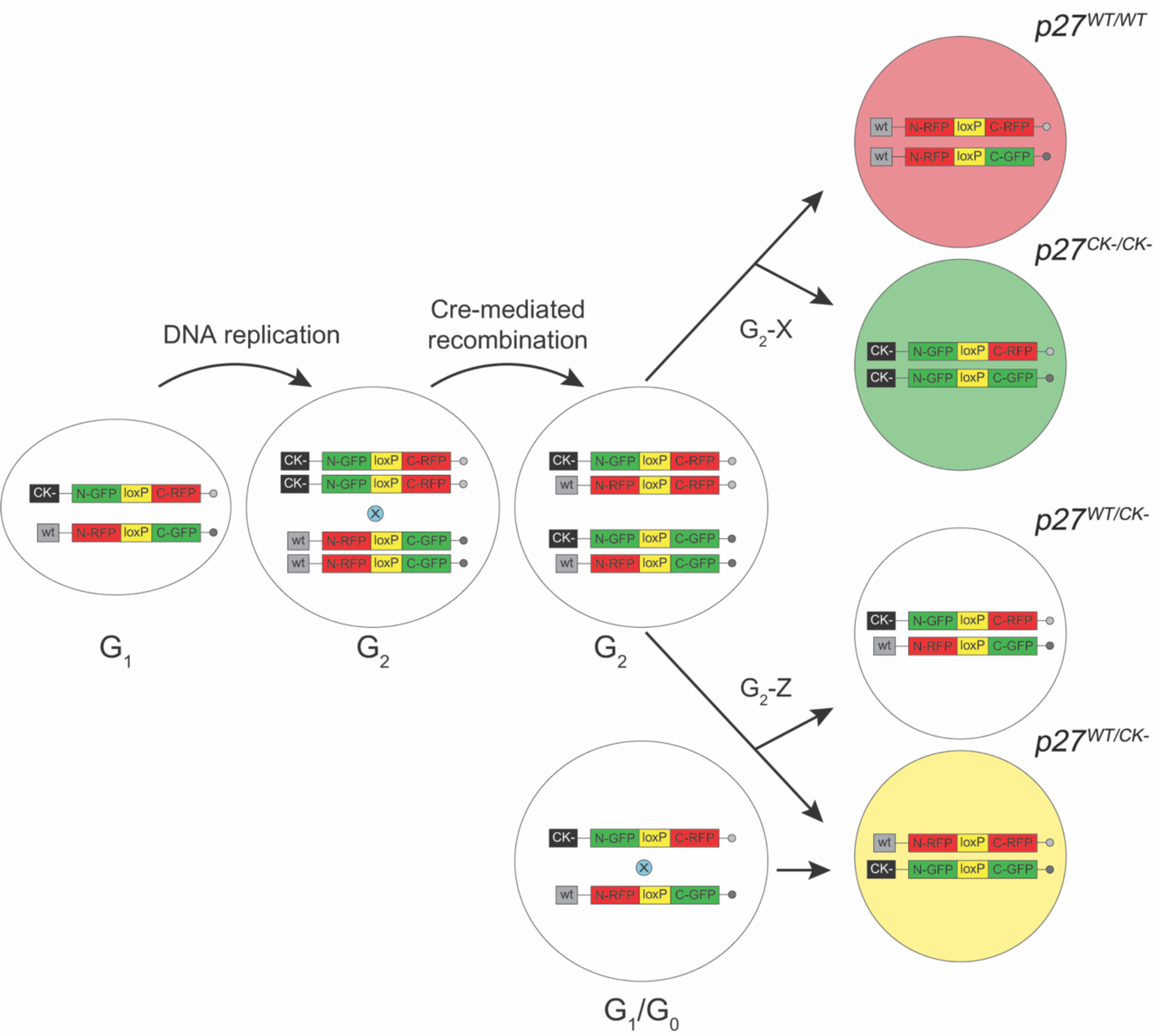
Diagram summarizing MADM outcomes. All experimental mice possess three transgenes: a ubiquitously-expressed Cre recombinase gene and two marker transgenes. Each marker transgene consists of partial N-or C-terminal coding sequences for GFP and RFP, reciprocally-arranged and interrupted by a single *loxP* site, on respective copies of chromosome 6. In a non-dividing cell (G_0_ or G_1_), Cre-catalyzed interchromosomal exchange results in functionally reconstituted fluorescent proteins in the same cell, which fluoresces yellow. However, in cycling cells S phase progression results in duplicated chromosomes which, upon functional recombination between homologous chromosomes in G_2_, produces differentially marked progeny. In the case of G_2_-X segregation, a pair of red and green cells is produced while G_2_-Z segregation results in colorless and yellow (double-labeled) cells. In the case of WT-MADM, all marked cells, regardless of color, are homozygous wild-type. However, in a heterozygous cell with a mutant *p27* gene (the CK-allele is depicted) linked to the chromosome carrying a GFP-RFP cassette (GR-MADM), Cre-recombination and G_2_-X segregation after mitosis yields a wild-type red cell (*p27^WT/WT^*) and a mutant green cell (*p27^CK-/CK-^*). G_2_-Z segregation of mitotic cells generates colorless and yellow cells, both heterozygous, while recombination taking place in either G_1_ or G_0_ generates double-labeled cells in all cases without altering genotype.

Adult mice older than 8-weeks were used for all experiments. Results from both male and female animals were pooled, since no apparent differences were noted between the sexes in preliminary experiments.

### Generation of MADM mice

Inherent in the MADM strategy is the unique and indelible labeling of the two daughter cells produced when a dividing cell undergoes interchromosomal recombination. In Fig 1, a cell containing the *MADM-6^GR^* and *MADM-6^RG^* transgenes, each on separate chromosomes, is depicted passing from G_1_ phase through S phase replication and into G_2_ phase. Prior to S phase recombination, the cell in G_1_ phase is unlabeled. Following DNA replication, Cre-induced rearrangement between *loxP* sites located on homologous chromosomes will result in restored GFP and RFP sequences in two of the four homologous chromosomes. Upon completion of mitosis, the labeled cells that result will depend on the mode of chromosomal segregation. In the case of X segregation, each cell will inherit one functional and one non-functional fluorescent protein gene, which upon expression will cause one cell to fluoresce red and the other green. Z segregation, on the other hand, will generate cells with either two reconstituted or two non-reconstituted transgenes (yellow and colorless, respectively). Importantly, unlike the situation when Cre activity takes place in G_2_ phase, recombination in non-mitotic cells during either G_0_ or G_1_ converts a colorless cell to a yellow cell that expresses both fluorescent proteins (Fig 1). Thus, in MADM mice the segregation within a cell of functionally recombined DsRed or GFP genes is linked to the generation of either a red or green cell, respectively.

If the above recombination event takes place on a wild-type background, the result will be marked cells that are phenotypically normal, a situation referred to as WT-MADM. This approach has been used successfully to fill restricted numbers of CECs with fluorescent label for cell shape analysis (5). GR-MADM, a variant strategy appropriate for single-cell phenotypic analysis, is made possible when one chromosome contains a mutation of interest distal (telomeric) to the knockin site of the MADM cassette (in this case, *Rosa26*). Because the red/green transgenes are linked to the wild-type/mutant alleles, respectively, through meiotic recombination, G_2_ recombination followed by X segregation (i.e., a G_2_-X event) will generate a green daughter cell homozygous for the mutation and a red sibling homozygous for the wild-type allele. G_2_-Z, G_1_ or G_0_ events, on the other hand, will produce cells that remain heterozygous and, because of their color (yellow) or lack thereof, are distinguishable from single color homozygous cells. In all cases, continued division of any one of the marked cells involved will thereafter lead to daughter cells of the same genotype. Finally, because the likelihood of recombination is relatively low at any one mitotic event, marked cells are generally rare and widely distributed, making the identification of such events more feasible (36, 38, 39).

In our experiments, implementation of WT-MADM was accomplished through a series of two crosses (40). First, homozygous *MADM-6^RG/RG^* and *Hprt^Cre/Cre^* mice were interbred to produce double heterozygotes. The latter were then mated with *MADM-6^GR/GR^* animals to generate Cre-positive mice (*MADM-6^GR/RG^*; *Hprt^WT/Cre^*) used for experiments and Cre-negative animals (*MADM-6^GR/RG^*; *Hprt^WT/WT^*), which provided a control. GR-MADM was carried out separately for the *p27^CK−^* and *p27*^−^ mutant strains (40). In both cases, *MADM-6^GR/GR^* mice were first bred to heterozygous mutant animals (*p27^+/^*^−^ or *p27^+/CK−^*) and double heterozygotes (*MADM-6^WT/GR^*; *p27^+/^*^−^ or *MADM-6^WT/GR^*; *p27 ^+/CK−^*) identified by genotyping. The latter were then backcrossed to wild-type C57Bl/6J mice to produce progeny that were screened by PCR for meiotic recombinants in which a mutant gene allele (*p27^CK−^* or *p27*^−^) was linked to the *MADM-6^GR^* transgene (designated *p27 MADM-6^WT/GR^*). Finally, *p27 MADM-6^WT/GR^* mice were crossed with *MADM-6^WT/RG^*; *Hprt^WT/Cre^* double heterozygotes (generated as above from *MADM-6^RG/RG^* and *Hprt^Cre/Cre^* animals) in order to produce *p27 MADM-6^GR/RG^*; *Hprt^WT/Cre^* experimental mice and *p27 MADM-6^GR/RG^*; *Hprt^WT/WT^* controls.

### Tissue preparation and immunostaining

Mice were euthanized using CO_2_ inhalation, followed by decapitation and pneumothorax. The eyes were then surgically isolated, punctured posteriorly with a 27-gauge needle and fixed overnight at 4°C in 2% paraformaldehyde in sodium acetate buffer, pH 6.0 (41). After several washes in phosphate-buffered saline, pH 7.3 (PBS), the globes were dissected to remove all non-corneal tissues and processed for reporter visualization and/or immunocytochemistry (see below). In most cases, corneas were kept intact for subsequent flat-mounting (5). Occasionally, they were cryoprotected with 20% sucrose overnight, frozen in Tissue-Tek O.C.T. compound (Sakura Finetek USA, Torrance, CA) and sectioned at 10µm using a cryomicrotome.

In order to visualize cell boundaries, corneas from wild-type mice and whole-animal mutants were labeled with an antibody to the tight junction-associated protein ZO-1. To suppress non-specific binding of the anti-mouse secondary antibody, tissues underwent a two-stage blocking reaction. In the first stage, they were incubated in normal donkey serum (NDS; Jackson ImmunoResearch Laboratories, West Grove, PA), diluted 1:10 in Tris-buffered saline (TBS), pH 7.3, with 1.0% bovine serum albumin (BSA) and 0.4% Triton X-100 (TBS-BSA-Triton X-100; 1 h; room temperature). Then, after several rinses in TBS corneas were blocked with goat anti-mouse IgG Fab fragments (Jackson ImmunoResearch; 0.13 mg/ml; 1 h; room temperature) prior to overnight incubation at 4°C in mouse anti-ZO-1 (Life Technologies, Carlsbad, CA; 1A12; 1:100). Following further buffer washes, tissues were stained for 2 h at room temperature with Alexa Fluor 488–conjugated donkey anti-mouse IgG (Life Technologies) before being rinsed and flat-mounted.

Intact corneas or corneal sections from MADM mice were blocked with NDS, then incubated overnight at 4°C in combined primary antibodies (chicken anti-green fluorescent protein [GFP]; Aves Labs, Tigard, OR; 1:500 and goat anti-c-Myc; Novus Biologicals, Littleton, CO; 1:200) diluted in TBS-BSA-Triton X-100. The c-Myc antibody was pre-absorbed with fixed wild-type corneal tissue pieces before use (see (42)). The following day, specimens were stained with fluorescein isothiocyanate (FITC)–conjugated anti-chicken IgG (1:200 dilution) and Alexa Fluor 555–conjugated anti-goat IgG (1:400 dilution) to enhance visualization of EGFP and c-Myc-DsRed2, respectively. After several buffer rinses, four radial incisions were made in each cornea to allow flattening of the tissue, which was then placed on a glass slide with the endothelium facing up and coverslipped using Vectashield fluorescence mounting medium (Vector Labs, Burlingame, CA) (32).

### Microscopy, image analysis and statistics

Corneal tissues were examined in either a Leica TCS SP2 or SP8 confocal laser scanning microscope equipped with standard excitation and emission filters (Leica, Heidelberg, Germany). Images from anti-ZO-1-stained endothelial monolayers were acquired using a 20X (NA=0.7) infinity-adjusted objective at zoom=3.0, while those from MADM-labeled tissues employed a 10X (NA=0.4) objective and zoom=1.0. Maximum z-stack projections were assembled from individual optical slices (1024 × 1024 resolution) taken at 1μm intervals. Under these conditions, 3-5 slices were sufficient to include the entire endothelial cell layer. Tonal quality of the resulting projection images was adjusted using the Levels tool in Adobe Photoshop CS6 (Adobe Systems, San Jose, CA) prior to further processing.

Evaluation of endothelial cell density and collection of morphometric parameters were carried out using Fiji software (43). Within a corneal flat-mount, four central and four peripheral sites (one each per tissue quadrant) were selected for image acquisition. Central areas were defined as those encompassing the inner 50% of corneal surface area, while peripheral regions were limited to those within 200 μm of the cornea-sclera junction. Approximately 0.25mm^2^ of tissue (3.6% of the total) was sampled in this way. Using a custom ImageJ1 macro designed by one of us (JB; available at http://www.biovoxxel.de/wp/wp-content/uploads/2019/05/Endothelial Cells Detection_Final.ijm), cell boundaries where identified by an algorithm involving various convolution filters and a Voronoi diagram (based on Dirichlet tessellation). The detected cell areas were used as input for further analysis. Cell density and area, the number of neighboring cells and circularity (a measure of cell shape) were obtained using the Extended Particle Analyzer and Neighbor Analysis functions of the ImageJ BioVoxxel Toolbox. For each parameter, the four central and peripheral measurements were averaged separately.

To compare patterns of cell proliferation and radial dispersion in mosaic corneas, panoramas encompassing the full expanse of labeled endothelia were assembled from multiple overlapping Z-stacks. In initial studies, this was accomplished using the Photomerge command of Photoshop, with individual single- and double-labeled cells counted manually in the composite images. In subsequent analyses, we made use of a macro (designed by JB) that incorporates Z-projection image stitching along with automatic detection and annotation of labeled cells. Twin spots (MADM-generated clones containing wild-type and mutant cells) were defined as groups of adjacent red and green cells located at least 200μm from the nearest single-labeled cell. They were identified using an algorithm that exploits cell proximity measurements to discover closely associated red and green cells within a circle of defined radius, followed by manual elimination of possible overlapping clones. A best fit was achieved in WT-MADM corneas with a radius of 82 pixels (248μm), which was then applied to mutant mosaic tissues. Analysis of the spatial arrangement of single-labeled cells within endothelia was implemented using a size and shape invariant modification of the density-based clustering algorithm DBSCAN for 2D images (44). In this unsupervised algorithm, dense regions (clusters) within an array of points are identified by the maximum distance between points (ε) and the minimum number of points (minimum density) in a cluster. Data in the present study were obtained using ε=45 pixels (136μm, cluster radius) and minimum density=4 (minimum cluster size). All of the processing and analytical tools described are available as partially individual plugins as part of the ImageJ BioVoxxel Toolbox (SSIDC Cluster Indicator) or as ImageJ1 macro code (complete processing pipeline for cluster and twin spot analysis from http://www.biovoxxel.de/wp/wp-content/uploads/2019/05/SSIDC_Cluster_And TwinSpot Analysis Final.ijm). Instructions for its use can be found under https://imagej.net/BioVoxxel Toolbox or in the respective macro code.

For whole-animal studies, one cornea per mouse was used to generate quantitative data. On the other hand, in reporting the MADM results we made use of both tissues of a single animal where possible. Our rationale for this is based on the assumption that the number and developmental timing of recombination events are random events and, thus, uncorrelated between eyes. The substantial amount of variability observed between endothelia of an individual mouse (e.g., green cell/red cell ratios differ by as much as 87%) confirmed this prediction. In such cases, both corneas of an animal were included in the analysis in order to document this variation and provide a fuller picture of the data.

All data were presented as means ± SEM. Statistical analysis was performed using Prism 8.0 software for Mac (GraphPad Software, San Diego, CA).

## Results

### The corneal endothelium of whole-organism *p27^CK-/CK-^* and *p27^−/−^* mutants are indistinguishable phenotypically

We first examined the effects of the *p27^CK^*^−^ and *p27*^−^ alleles on endothelial cell density in whole-organism mutants. Representative images of tissue flat-mounts from homozygous wild-type and mutant corneas are displayed in Fig 2A-2F. Compared with wild-type monolayers, central regions of both *p27^CK^*^−^*^/CK^*^−^ and *p27*^−/−^ endothelia exhibit density enhancements of approximately 60% (Fig 2G). A very similar quantitative relationship is also seen for peripheral regions, although the latter maintain consistently lower cell densities relative to central areas in all genotypes (Fig 2G).

**Figure 2.**
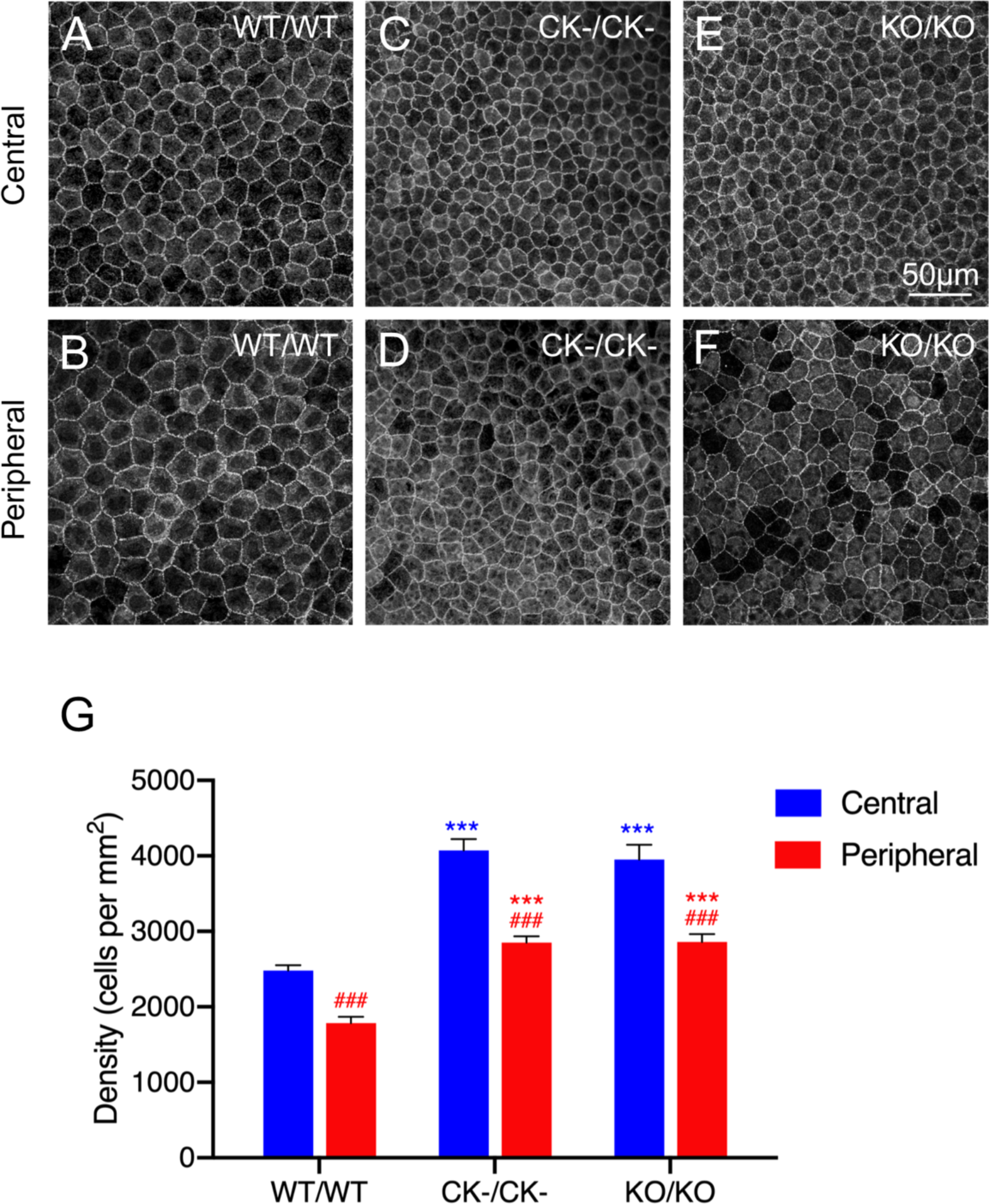
Analysis of cell density in endothelial monolayers from organism-wide *p27^+/+^*, *p27^CK−/CK−^* and *p27^−/−^* mice (WT/WT, CK-/CK- and KO/KO, respectively). (A-F) Apical cell boundaries revealed by anti-ZO-1 labeling. Compared with wild-type controls (A and B), increases in cell density are seen in both central and peripheral regions of *p27^CK−/CK−^* (C and D) and *p27^−/−^* (E and F) corneas. For each genotype, peripheral regions exhibit consistent decreases in density relative to corresponding central regions. (G) Quantitation of cell density. Data represent means ± SEM (n = 11 [wild-type] or 6 [mutants]). Ordinary two-way ANOVA followed by Tukey’s HSD test was performed. *** and *** indicate p<0.001 by comparison to wild-type, while ### indicates p<0.001 by comparison to central regions.

One consequence of the increased cell densities in mutant endothelia is corresponding decreases in average cell size, estimated as apical face area (S1A Fig). When the data are plotted as size distributions, these decreases are seen to be the result of regionally distinct transformations. In the case of central areas, there is a shift in the peak of mutant cell sizes to smaller values, with relatively little change in peak widths (S1B-S1D Fig). On the other hand, for peripheral regions the peak shift is accompanied by a more compressed size distribution, as well as a greater overlap with central values (S1B-S1D Fig).

To investigate whether the increased cell density seen in *p27^CK-/CK-^* and *p27*^−/−^ mutants has an impact on the apical cell surface geometry, analyses of cell shape and nearest neighbors were carried out. Our results show that, while there is a small but significant increase in circularity (regularity of shape) for cells in central regions compared to those in peripheral regions across all genotypes, little difference is seen between the respective regional populations of wild-type and mutant endothelial cells (S2A Fig). Similarly, neighbor analysis indicates that the distribution of neighbors per cell, and thus the arrangement of cells within monolayers, is unchanged by gene knock-out or mutation (S2B-S2D Fig).

### MADM allows analysis of single cells generated during endothelial development

To investigate *p27* gene function at the level of individual cells, mosaic analysis was carried out. Initially, corneas from WT-MADM mice were examined to assess cell labeling and establish a baseline for mutant studies. When these tissues are viewed in cross-section, rare examples of labeled cells can be localized within the endothelial cell layer (arrowheads in Fig 3A). Similar specimens prepared as flat-mounts allow all GFP^+^ and RFP^+^ cells to be seen within the context of the monolayer (Fig 3B). While most are double-labeled (yellow), in every cornea variable numbers of single-labeled (green and red) cells are produced as a result of Cre-mediated G_2_X (mitotic recombination) events. These occur singly but are often components of discrete groups of cells in which one red cell or cluster lies adjacent to a green cell or cluster (Fig 3B, boxed areas). Since such groups are generally limited in number and widely separated within monolayers, they likely represent the progeny of single (interchromosomal) recombination events, referred to as twin spots, (40, 45). To explore this further, we determined the number and size of single-labeled cell groups using a proximity-based algorithm (see Materials and Methods). Our results show that, for WT-MADM tissues, an average of only about four such groups are observed per cornea (S3A Fig). Furthermore, the majority (83%) of these are of relatively small size (two to four cells each), suggesting that they represent independent clones, rather than the merger of cells from more than one MADM event (Fig 3C). Importantly, when considered both on the basis of whole corneas and individual twin spots green and red cells are generated in approximately equal numbers, indicating that expression of fluorescence markers does not differentially affect the expansion potential of wild-type sibling cells (Fig 3D).

**Figure 3.**
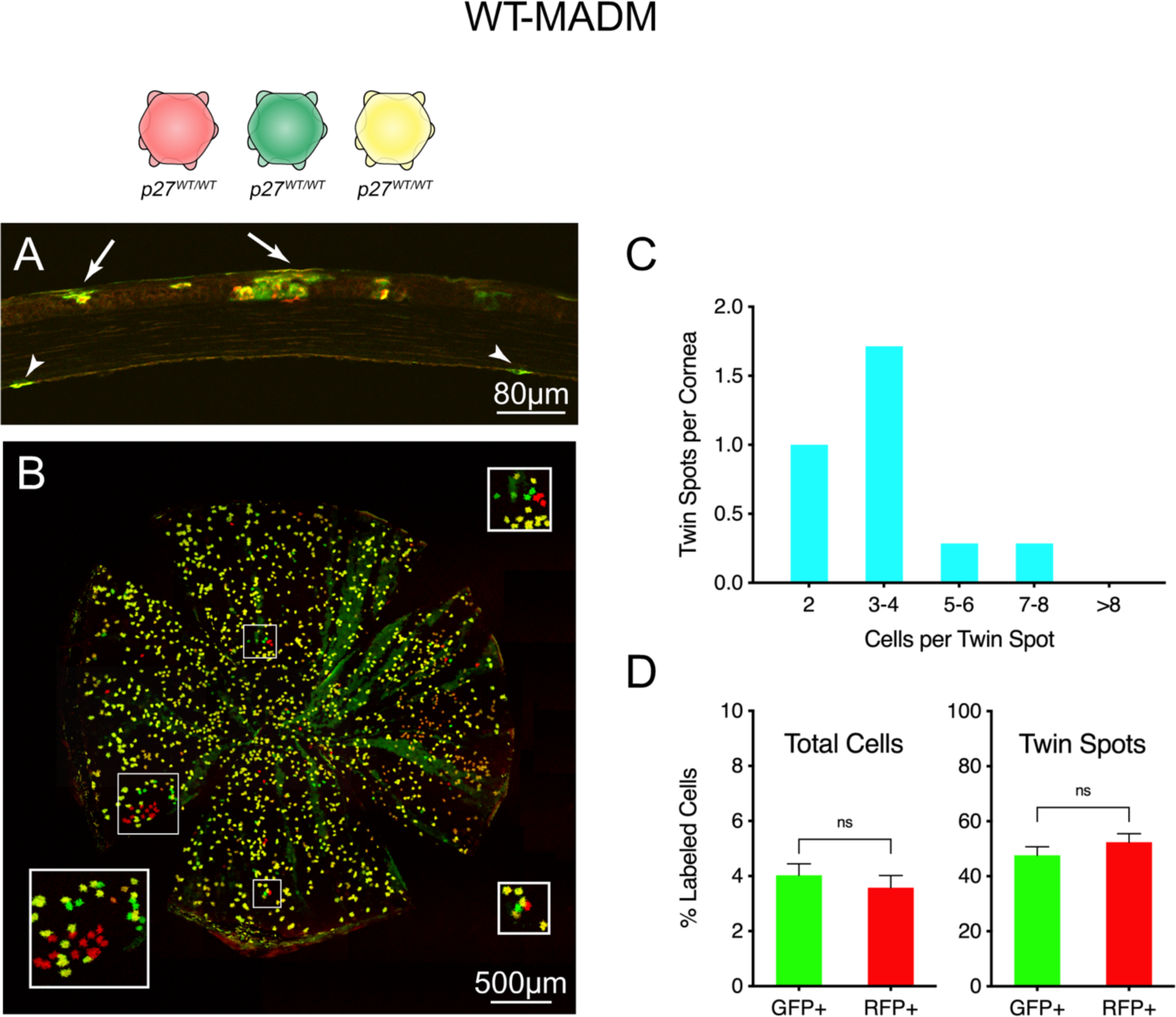
Visualization and analysis of single- and double-labeled cells in WT-MADM corneas. (A) Tissue section. In this full-thickness cross-section, the anterior corneal surface faces upward. Arrows point to groups of labeled cells within the multilayered epithelium, while arrowheads indicate two stained CECs within the endothelial monolayer. (B) Tissue whole-mount. Single- (red and green) and double-labeled (yellow) endothelial cells (all *p27^+/+^*) are confined to a thin tissue layer that is generally well-separated optically from fluorescent protein-expressing epithelial and stromal cells. In some places, however, moderate undulation of the flat-mounted tissue leads to bleed through from the epithelial layer (green radial streaks). Boxed areas indicate regions of interest (ROIs) containing groups of single-labeled cells resulting from G_2_X recombination events. Insets show boxed areas at 2X magnification. (C) Distribution of twin spot sizes (see Materials and Methods for a description of how twin spots were identified). The majority (83%) of twin spots contain two to four cells. (D) Relative proportion of single-labeled GFP^+^- and RFP^+^-positive cells in whole corneas and twin spots. Approximately equal percentages of MADM-generated red and green cells are observed, whether the data include the total number of single-labeled cells in corneas and are expressed relative to all labeled cells (red, green and yellow; left) or only the subset of single-labeled cells in individual twin spots (right). Values plotted in (C) result from analysis of 23 twin spots from seven corneas. Data in (D) represent means ± SEM (n = 5). Unpaired t-test was performed. ns indicates not significant.

### Endothelial cells expressing *p27^CK-^ and p27*^−^ alleles exhibit divergent proliferative and migration phenotypes

GR-MADM was performed in order to compare wild-type and mutant cells within the same endothelial monolayer. Examination of whole-mounted corneas from the *p27^CK^*^−^ and *p27*^−^ mosaic strains indicates that there are substantially more single-labeled cells present than in WT-MADM, with noticeably greater numbers of green cells than red cells (Fig 4A and 4B). To represent this increase of mutant cells relative to wild-type quantitatively, we made use of the green/red cell ratio as a metric (40). Our analysis of the entire population of single-labeled endothelial cells in *p27^CK^*^−^ mosaic corneas indicates that cells expressing a non-functional CDK inhibitor generate almost three times more progeny than those with wild-type p27 (Fig 4C). Surprisingly, *p27*^−^ mosaics, in which there is complete loss of CDK inhibitor expression, exhibit a further two-fold increase in the green-to red-cell ratio, resulting in an overall six-fold expansion of mutant to wild-type cells (Fig 4C).

**Figure 4.**
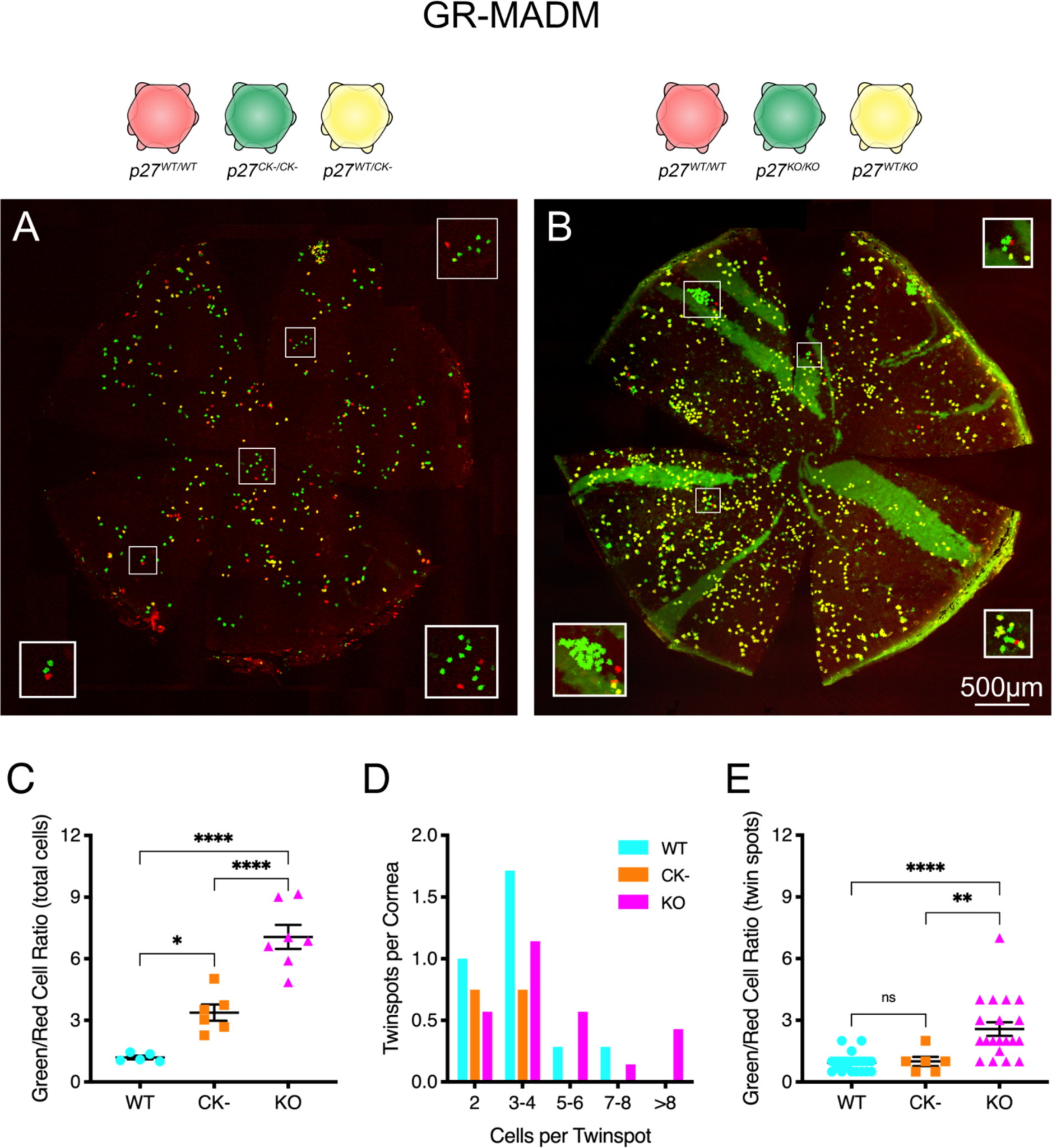
MADM analysis of homozygous *p27^CK−^*- and *p27^−^*-expressing CECs. (A and B) Whole-mounts of GR-MADM corneas (CK- and KO, respectively). Compared with WT-MADM tissues, increased numbers of single-labeled cells characterize corneas of both mutant genotypes. Boxed areas (ROIs) isolate groups of red and green cells, highlighting the enhanced numbers of mutant (GFP^+^) cells relative to wild-type (RFP^+^) cells in these areas. (C) Quantitative expansion of mutant versus wild-type cells. Each point represents the green-to-red cell ratio for all single-labeled cells of an individual cornea, with means for different genotypes indicated by horizontal lines. Roughly three- and six-fold increases in the ratio of mutant-to-wild-type cells is apparent for the *p27^CK−^* (CK-) and *p27^−/−^* (KO) mosaics, respectively. (D) Histogram of twin spot sizes for GR-MADM mutants, compared to WT-MADAM tissues (data replotted from Fig 3C). The few twin spots detected in *p27^CK−^* mosaic corneas are all relatively small. By contrast, the frequency distribution of these red and green cell-containing groups is shifted to larger sizes in *p27^−/−^* mosaics. (E) Expansion of mutant cells relative to wild-type cells in identifiable twin spots. Each point represents the green-to-red cell ratio for a single twin spot. Averaged data are represented as means ± SEM. In (C), n = 5 (WT), 6 (CK-) and 7 (KO). In (E), n = 23 (WT), 6 (CK-) and 20 (KO). Statistics was conducted by ordinary two-way ANOVA followed by Tukey’s HSD test. *, **, and **** indicate p<0.05, p<0.005 and p<0.0001, respectively. ns indicates not significant. Values plotted in (D) result from analysis of six twin spots from four corneas (*p27^CK−^* mosaics) and 20 twin spots from seven corneas (*p27^−/−^* mosaics).

To determine whether these mutation-specific differences are due to changes in the proliferative potential of individual cells, we performed twin spot analysis. Using the same criteria employed for WT-MADM corneas, twin spot number and size are seen to differ significantly between the two mutants (Fig 4D). In the case of *p27^CK^*^−^ mosaic corneas, coherent groups of GFP+ and RFP+ cells are relatively rare and, similar to wild-type tissues, contain only two to four cells each. By contrast, a shift in the distribution of *p27*^−^ mosaic twin spots to larger sizes results in more single-labeled cells being found in these arrays. Importantly, twin spots in the latter corneas have substantially higher green/red cell ratios than those from either *p27^CK^*^−^ mosaics or *p27^+^* tissues, indicating an increased proliferative potential for cells of this genotype (Fig. 4E).

While endothelia in both *p27^CK^*^−^ and *p27*^−^ mosaics have elevated numbers of mutant green cells, due to increased recombination events and enhanced cell division, the resulting cells are distributed very differently within their respective tissues. Fig 5 shows representative regions, from both wild-type and mutant mosaic corneas, where red and green cells are relatively abundant. Single-labeled cells in twin spots of WT-MADM tissues are generally located within one or a few cell diameters of one another (Fig 5A-A’’). Because of the rarity of recombination events, this arrangement results in small discrete groups of clustered cells. In *p27^CK^*^−^ mosaics, on the other hand, individual green and red cells appear scattered within the corneal monolayer, with extensive areas of unlabeled cells interposed between them (Fig 5B-B”). The fact that obvious groups of single-labeled cells are less evident, suggests a greater degree of mobility of cells away from sites of MADM-generation in these corneas. Interestingly, clusters of green cells in *p27*^−^ mosaics appear to be even more delimited than those of wild-type tissues. Examples of close-packed GFP+ cells, along with associated RFP+ cells, are depicted in Fig 5C-C”. In these clusters, adjacent mutant cells often appear to directly touch one another. The highly restricted distribution of single-labeled cells, coupled with the expanded generation of mutant *p27*^−^ mutant cells characteristic of these mosaics, often leads to large clusters of GFP+ cells (Fig 5C).

**Figure 5.**
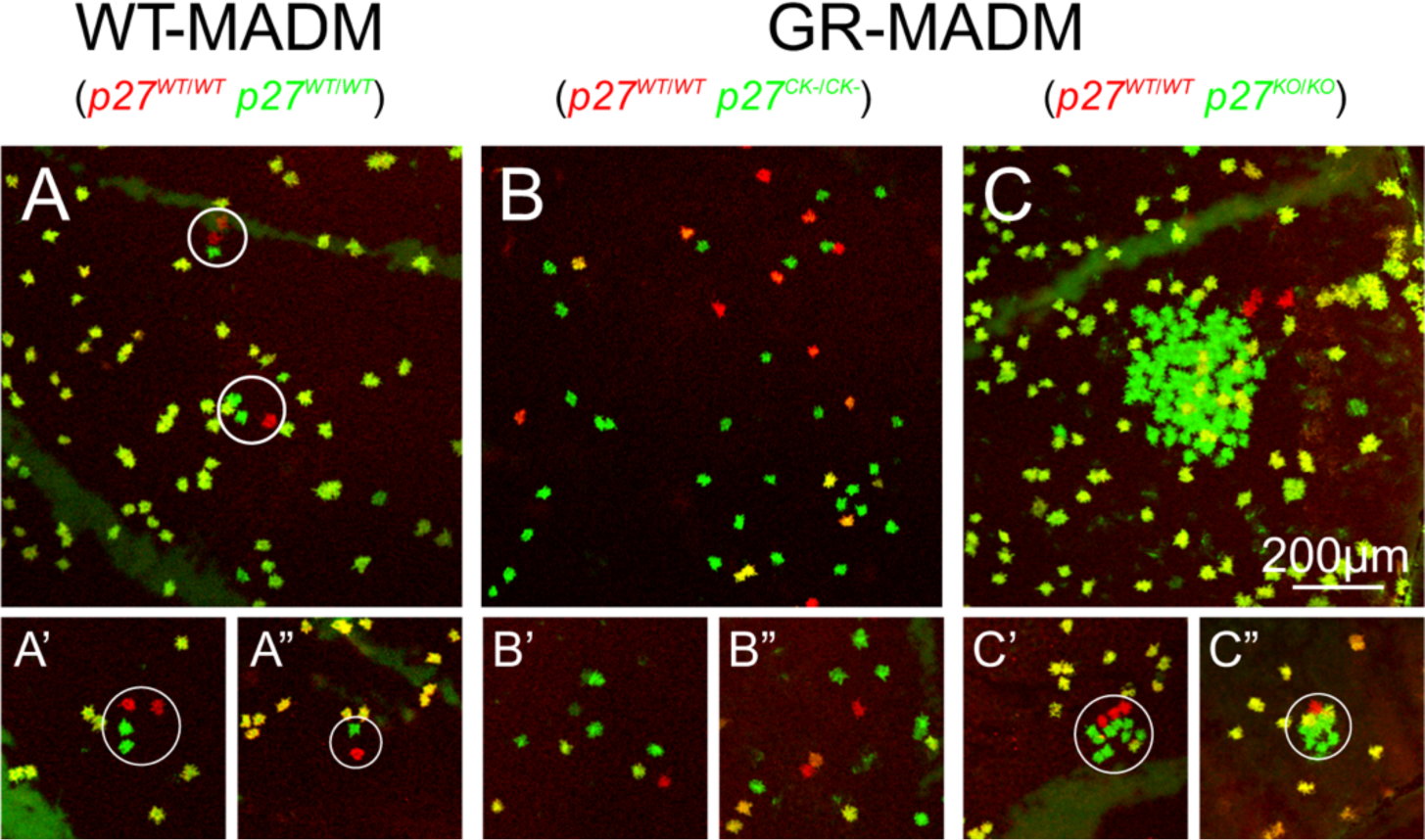
Relative dispersion of MADM-generated single-labeled cells in wild-type and mutant mosaic endothelia. (A-A”) WT-MADM. In twin spots (circled), red and green cells (both *p27^+/+^*) directly abut one another or are separated by, at most, one or two cell diameters. (B-B”) *p27^CK−^* GR-MADM. Note that GFP^+^- and RFP^+^-positive cells (mutant and wild-type, respectively) are more scattered within monolayers, with variable regions of unlabeled cells separating them. None of the cells in these ROIs belong to twin spots. (C-C”) *p27^KO^* GR-MADM. Compared with the other two genotypes, green cells (*p27^−/−^*) are packed together with little intervening space and oftentimes appear to be in direct contact with one another and occasional yellow cells (*p27^+/−^*). Because of its large size, the group of single-labeled cells in (C) did not qualify as a single twin spot and, thus, is not circled.

Whole-endothelium plots of single-labeled cells reveal global patterns of cell distribution that mirror the differences seen in dispersion of GFP+ and RFP+ cells from local sites of G_2_-X recombination. In WT-MADM corneas, red and green cells are observed mainly in small clusters, although they also exist singly (Fig 6A). While *p27^CK^*^−^-expressing cells are evenly distributed across endothelial monolayers, *p27*^−/−^ cells are confined to relatively few highly coherent clusters (Fig 6B and 6C). To quantify these patterns, we used the density-based algorithm DBSCAN to perform cluster analysis. Fig 6D illustrates that, compared with *p27*^+/+^-expressing cells in WT-MADM tissues, the percentages of *p27^CK^*^−^*^/CK^*^−^ and *p27*^−/−^ cells in clusters is two-and four-fold greater, respectively.

**Figure 6.**
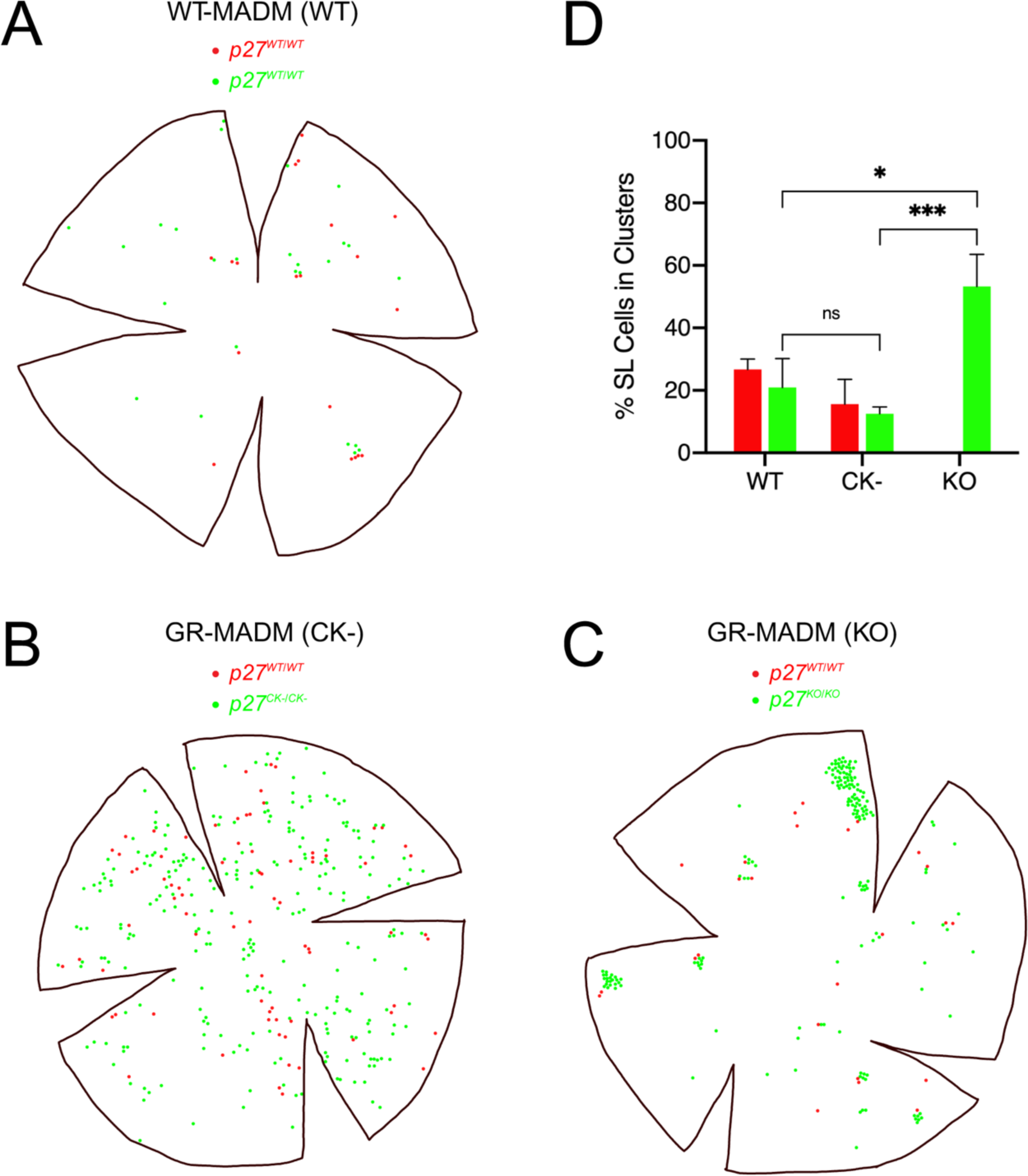
Analysis of cell distribution patterns in mosaic wild-type and mutant corneas. (A-C) Representative plots of GFP^+^ and RFP^+^ cells in WT-MADM (A), *p27^CK−^* GR-MADM (B) and *p27^KO^* GR-MADM (C) tissues. In (A), red and green cells (both *p27^+/+^*) appear in small clusters, as well as distributed as single cells within corneas. Many more MADM-generated cells are apparent in *p27^CK−^* and *p27^KO^* mosaics. However, the distribution patterns of GFP^+^ cells differ greatly between the two mutant mosaics. In *p27 ^CK−^* GR-MADM tissues (B), homozygous mutant (green) cells are evenly dispersed within endothelial monolayers. By contrast, GFP^+^ cells in *p27^KO^* GR-MADM corneas (C) appear mainly in highly coherent clusters. (D) Cluster analysis. Compared with single-labeled (SL) GFP^+^ cells from wild-type and CK-mosaics, MADM-generated mutant cells in *p27^KO^* GR-MADM corneas are two- and four-fold more likely to be found in clusters. Bars represent means ± SEM (n = 4 [red cells] and 3 [green cells]) for WT, 4 [red cells] and 6 [green cells] for CK- and 7 [green cells] for KO). There were no clusters of *p27^+/+^* (red) cells in *p27^KO^* mosaics. Statistical analysis was carried out using ordinary two-way ANOVA followed by Sidak’s multiple comparisons test. * and *** indicate p<0.005 and p<0.001, respectively. ns indicates not significant.

## Discussion

p27 is a pivotal factor regulating cell cycle withdrawal in many lineages of developing organisms, including those of the corneal endothelium. In this study, we compared proliferation and migration phenotypes of murine CECs expressing either partial (*p27^CK^*^-^) or full (*p27^−^*) loss-of-function alleles of the *cdk1b* gene. Whole animal mutants, in which all cells are affected, exhibited moderate (∼0.6-fold) endothelial hyperplasia, with no quantitative difference between the two variant strains with respect to the final cell density achieved. On the other hand, in endothelia with sporadically-generated homozygous mutant and wild-type cells there was a much greater increase in mutant cell number expansion, as well as a two-fold difference between mice with the *p27^CK^*^-^ and *p27^−^* alleles.

The approximately six-fold enhancement in MADM-generated mutant versus wild-type cells is similar to that previously reported for cerebellar granule cells by Muzumdar et al. (40). As discussed by these authors, proliferation of mutant cells may be less restricted in mosaic mice because they constitute a relatively small proportion of the tissues in which they are produced. Thus, actively dividing cells would be expected to encounter less competition from surrounding wild-type cells than in the situation where all cells are mutant. Furthermore, because a smaller portion of the tissue is engaged in cell expansion, there would be less likelihood of activating inhibitory mechanisms regulating global tissue size.

An unexpected result from our study concerns the two-fold difference in the number of mutant cells, compared to wild-type, in mosaics expressing the *p27^CK^*^-^ and *p27^−^* alleles. p27 is a regulatory factor whose only direct influence on cell cycle progression is believed to be as an G_1_ cyclin-CDK inhibitor (25). Furthermore, previous studies have shown that this function is completely eliminated by mutations that alter four amino acids involved in cyclin and CDK binding (30, 35). For these reasons, it was predicted that homozygous expression of the *p27^CK^*^-^ allele would produce a phenotype equivalent to that of the *p27*-null allele. The fact that CEC proliferation is actually further enhanced in cells harboring two knockout alleles indicates that p27 negatively regulates cell cycle progression through an additional mechanism that is unrelated its role as a cyclin-CDK inhibitor.

One way that p27 could be acting in a non-canonical fashion is through effects on mitogenic signaling pathways that regulate cell cycle progression. For example, Fabris et al. (34) have recently shown using mouse embryo fibroblasts (MEFs) that cytoplasmic p27 regulates MAPK (ERK1/2) activation, and consequent early G_1_-S transit, through its effects on microtubule stability. This effect was shown to be dependent on the interaction of p27 with the microtubule-destabilizing protein stathmin, since increased proliferation in *p27*-null mice was reverted by concomitant deletion of the stathmin gene (33). Unlike the binding sites for CDKs which are located within the N-terminal portion of p27, the interaction with stathmin had been previously mapped to the C-terminal part of the protein (46). Interestingly, in these experiments the augmentation of cell division was due to faster cell cycle reentry whereas, using MADM, Muzumdar et al. (40) have demonstrated that deletion of *p27* results in extra cell cycles.

An alternative way in which p27 could be influencing CEC proliferation is through the small GTPase RhoA. Cytoplasmic p27 has been shown to inhibit Rho signaling by blocking the binding and activation of RhoA by Rho-GEFs (47). Coupled with the established involvement of activated Rho and its downstream effectors ROCK I and II in mitogen-stimulated G_1_-S phase progression, p27 could be functioning as an indirect modulator of cell division (48-50). In NIH 3T3 fibroblasts, where conditional activation of a ROCK-estrogen receptor fusion protein is sufficient to stimulate G_1_/S cell cycle progression, activated ROCK acts by upregulating cyclin D1 and cyclin A expression through Ras/MAPK and LIMK activation, respectively (51). Therefore, p27 could be exerting anti-proliferative effects both as a cyclin-CDK inhibitor in the nucleus and as a modulator of Rho/ROCK signaling in the cytoplasm.

While a role for stathmin in CEC proliferation has not been investigated, there is ample evidence for the involvement of ROCK in cell division of corneal endothelium. Initial experiments by Okumura et al. (52) using cultured monkey CECs demonstrated that the specific ROCK inhibitor Y-27632 promoted cell adhesion, inhibited apoptosis and increased the frequency of proliferating cells. Further studies have shown that these proliferative effects are due to coordinate upregulation of cyclin D and downregulation of p27 via enhanced phosphatidylinositol 3-kinase (PI-3-K)-Akt signaling (53). Evidence that active ROCK may normally be restraining cell division in adult CECs has been reproduced in several species, both in vitro and in vivo (54-57). In contrast to these studies, essentially opposite results of RhoA and ROCK inhibitors on endothelial cell proliferation have been reported by Zhu et al. (58). When siRNA against the p120 catenin was administered to human CEC cultures, activating trafficking of this adherens junction component to the nucleus, BrdU incorporation was stimulated in parallel with increased RhoA-ROCK signaling. Furthermore, this BrdU incorporation was abolished by inhibitors targeting either RhoA (CT-04) or ROCK (Y27632). While these data appear to be in conflict with one another, there is evidence that the direction of cell division upon Rho/ROCK signaling can be cell type- or context-specific (50). Thus, these studies suggest that, under different conditions, ROCK can be either an activator or an inhibitor of cell division.

In addition to proliferative changes, our experiments reveal major differences in the distribution of MADM-generated wild-type and mutant CECs within developing endothelial cell monolayers. Because pairs of red and green cells that arise by G_2_-X recombination initially lie directly adjacent to one another, the extent of dispersion of these cells and their progeny is an indication of their radial migration. The fact that most labeled *p27^+/+^* cells generated in WT-MADM tissues remain in relatively close proximity to one another indicates that CECs normally exhibit limited motility within the developing monolayer. Consequently, the greater dispersion of *p27^CK^*^−^-expressing cells suggests enhanced migration, while the more highly clustered arrangement of *p27^-^*^-^-expressing cells indicates more restricted migration.

Previously, it’s been demonstrated that cultured p27-deficient MEFs exhibit impaired motility in combination with an increased number and size of actin stress fibers and focal adhesions (47, 59, 60). This was shown to be the result of increased Rho-GTP levels due to loss of p27 and its ability to directly bind and inhibit Rho activation by guanine–nucleotide exchange factors (GEFs) (47). Consequently, Besson et al. (47) have proposed that p27 normally acts as a positive regulator of cell migration by titrating the Rho signal and allowing for cell movement through a balance of Rac and Rho activities. Supporting the importance of this pathway in developing tissues, regulation of RhoA by p27 has been found to be important for proper migration of neuronal progenitor cells in the subventricular cortex of developing mice (61, 62). In contrast to this work, Baldassarre et al. (46) have presented evidence that, in cultured sarcoma cells, p27 is normally a negative regulator of migration, while its binding to stathmin is able to counteract this migration inhibition.

The highly restricted distribution of *p27*-null cells seen in our MADM experiments is consistent with reduced motility due to excessive Rho activity. Conversely, the fact that dispersion of homozygous *p27^CK^*^-^-expressing CECs is actually increased may be related to expression of the inhibitor. Besson et al. (47) have previously noted that MEFs from *p27^CK^*^-^ knock-in mice produce levels of p27 above those seen in wild-type controls, most likely because cyclin and CDK binding is important for turnover of the protein (30, 35, 63). If a similar increase takes place in *p27^CK^*^-^ mosaic corneas, this could account for the apparent enhancement in migration by mutant CECs that we see since the migration-promoting activity of p27 lies in the C-terminal half of the protein, which would be intact in *p27^CK^*^-/*CK*-^ mutants (59).

Overall, our results highlight unexpected complexity in the way that the cell cycle effector protein p27 acts during construction of the corneal endothelium. The fact that its mutation has profound consequences for both cell proliferation and migration underscores the intimate link between these two functions in cells and the role of p27 in their integration. Finally, our studies further demonstrate the advantages of single cell mutational analysis for unravelling protein function within the context of developing mammalian tissues.

## Acknowledgements

This research was supported by grants from the National Institutes of Health (R15 EY17997) and the ETSU Research Development Committee (to DMD). It was also supported by NIH grant C06RR0306551. We wish to thank Dr. Simon Hippenmeyer of the Institute of Science and Technology Austria for helpful advice on the early stages of this study.

## Author Contributions

Conceived and designed the experiments: DMD TAH. Performed the experiments: PDM HR. Developed new analytical methods: JB. Analyzed the data: DJH DMD. Wrote the paper: DMD TAH.

## References

1. Kreutziger GO. Lateral membrane morphology and gap junction structure in rabbit corneal endothelium. Exp Eye Res. 1976; 23:285–93. [PMID: 976372]

2. Hirsch M, Renard G, Faure J-P, Pouliquen Y. Study of the ultrastructure of the rabbit corneal endothelium by the freeze-fracture technique: apical and lateral junctions. Exp Eye Res. 1977 Sep; 25(3):277–88. [PMID: 590370]

3. Ringvold A, Davanger M, Olsen EG. On the spatial organization of the cornea endothelium. Acta Ophthalmol. 1984 Dec; 62(6):911–8. [PMID: 6524316].

4. Forest F, Thuret G, DuMollard JM, Peoc’h M, Perrache C, He Z. Optimization of immunostaining on flat mounted human corneas. Mol Vis. 2015 Dec 30; 21:1345–1356. [PMID: 26788027]

5. Harrison TA, He Z, Boggs K, Thuret G, Liu HX, Defoe DM. Corneal endothelial cells possess an elaborate multipolar shape to maximize the basolateral to apical membrane area. Mol Vis. 2016 Jan 16;22:31–9. [PMID: 27081293].

6. He Z, Forest F, Gain P, Rageade D, Bernard A, Acquart S, et al. 3D map of the human corneal endothelial cell. Sci Rep. 2016; Jul 6;6:29047. doi: 10.1038/srep29047. [PMID: 27381832]

7. Bonanno JA. Molecular mechanisms underlying the corneal endothelial pump. Exp Eye Res. 2012;95:2–7. doi: 10.1016/j.exer.2011.06.004. [PMID: 21693119].

8. Wilson RS, Roper-Hall MJ. Effect of age on the endothelial cell count in the normal eye. Br J Ophthalmol. 1982 Aug;66(8):513–5. [PMID: 7104267].

9. Murphy C, Alvarado J, Juster R, Maglio M. Prenatal and postnatal cellularity of the human corneal endothelium. A quantitative histologic study. Invest Ophthalmol Vis Sci 1984 Mar;25(3):312–22. [PMID: 6698749].

10. Joyce NC. Proliferative capacity of the corneal endothelium. Prog Retin Eye Res. 2003 May;22(3):359–89. [PMID: 12852491].

11. Laing RA, Sanstrom MM, Berrospi AR, Leibowitz HM. Changes in the corneal endothelium as a function of age. Exp Eye Res. 1976 Jun;22(6):587–94. [PMID: 776638].

12. Laule A, Cable MK, Hoffman CE, Hanna C. Endothelial cell population changes of human cornea during life. Arch Ophthalmol. 1978 Nov;96(11):2031–5. [PMID: 718491].

13. Sherrard ES. The corneal endothelium in vivo: its response to mild trauma. Exp Eye Res. 1976 Apr;22(4):347–57. [PMID: 954871].

14. Yee RW, Matsuda M, Schultz RO, Edelhauser HF. Changes in the normal corneal endothelial cellular pattern as a function of age. Curr Eye Res. 1985 Jun;4(6):671–8. [PMID: 4028790].

15. Ikebe H, Takamatsu T, Itoi M, Fujita S. Age-dependent changes in nuclear DNA content and cell size of presumably normal human corneal endothelium. Exp Eye Res. 1986 Aug;43(2):251–8. [PMID: 3758224].

16. Losick VP, Jun AS, Spradling AC. Wound-induced polyploidization: regulation by Hippo and JNK signaling and conservation in mammals. PLoS One 2016 Mar 9;11(3):e0151251. doi: 10.1371/journal.pone.0151251. [PMID: 26958853].

17. Mishima S. Clinical investigations on the corneal endothelium-XXXVIII Edward Jackson Memorial Lecture. Am J Ophthalmol. 1982 Jan;93(1):1–29. [PMID: 6801985].

18. Schmedt T, Silva MM, Ziaei A, Jurkunas U. Molecular bases of corneal endothelial dystrophies. Exp Eye Res. 2012 Feb;95(1):24–34. doi: 10.1016/j.exer.2011.08.002. [PMID: 21855542].

19. Tan DT, Dart JK, Holland EJ, Kinoshita S. Corneal transplantation. Lancet. 2012 May 5;379(9827):1749-61. doi: 10.1016/S0140-6736(12)60437-1. [PMID: 22559901].

20. Tuft SJ, Coster DJ. The corneal endothelium. Eye. 1990;4(Pt 3):389–424. doi: 10.1038/eye.1990.53. [PMID: 2209904].

21. Joyce NC, Harris DL, Zieske JD. Mitotic inhibition of corneal endothelium in neonatal rats. Invest Ophthalmol Vis Sci. 1998 Dec;39(13):2572–83. [PMID: 9856767].

22. Gordon SR. Changes in extracellular matrix proteins and actin during corneal endothelial growth. Invest Ophthalmol Vis Sci 1990 Jan;31(1):94–101. [PMID: 2404898].

23. Joyce NC, Harris DL, Mello DM. Mechanisms of mitotic inhibition in corneal endothelium: contact inhibition and TGF-beta2. Invest Ophthalmol Vis Sci. 2002 Jul;43(7):2152–9. [PMID: 12091410].

24. Yoshida K, Kase S, Nakayama K, Nagahama H, Harada T, Ikeda H, et al. Involvement of p27^KIP1^ in the proliferation of the developing corneal endothelium. Invest Ophthalmol Vis Sci. 2004 Jul;45(7):2163–7. [PMID: 15223790].

25. Sherr CJ, Roberts JM. CDK inhibitors: positive and negative regulators of G_1_-phase progression. Genes Dev. 1999 Jun 15;13(12):1501–12. [PMID: 10385618].

26. Russo AA, Jeffrey PD, Patten AK, Massagué J, Pavletich NP. Nature 1996 Jul 25;382(6589):325-31. doi: 10.1038/382325a0. [PMID: 8684460].

27. Fero ML, Rivkin M, Tasch M, Porter P, Carow CE, Firpo E, et al. A syndrome of multiorgan hyperplasia with features of gigantism, tumorigenesis, and female sterility in p27*^Kip1^*-deficient mice. Cell. 1996 May 31;85(5):733–44. [PMID: 8646781].

28. Kiyokawa H, Kineman RD, Manova-Todorova KO, Soares VC, Hoffman ES, Ono M, et al. Enhanced growth of mice lacking the cyclin-dependent kinase inhibitor function of p27*^Kip1^*. Cell. 1996 May 31;85(5):721–32. [PMID: 8646780].

29. Nakayama K, Ishida N, Shirane M, Inomata A, Inoue T, Shishido N, et al. Mice lacking p27*^Kip1^* display increased body size, multiple organ hyperplasia, retinal dysplasia, and pituitary tumors. Cell. 1996 May 31;85(5):707–20. [PMID: 8646779].

30. Besson A, Gurian-West M, Chen X, Kelly-Spratt KS, Kemp CJ, Roberts JM. A pathway in quiescent cells that controls p27^Kip1^ stability, subcellular localization, and tumor suppression. Genes Dev 2006 Jan 1;20(1):47–64. doi: 10.1101/gad.1384406. [PMID: 16391232].

31. Besson A, Hwang HC, Cicero S, Donovan SL, Gurian-West M, Johnson D, Clurman BE, Dyer MA, Roberts JM. Discovery of an oncogenic activity in p27Kip1 that causes stem cell expansion and a multiple tumor phenotype. Genes Dev 2007 Jan 1;21(1):1731–1746. doi: 10.1101/gad.1556607. [PMID: 17626791].

32. Defoe DM, Adams LB, Sun J, Wisecarver SN, Levine EM. Defects in retinal pigment epithelium cell proliferation and retinal attachment in mutant mice with p27^Kip1^ ablation. Mol Vis. 2007 Feb 27;13:273–86. [PMID: 17356514].

33. Berton S, Pellizzari I, Fabris L, D’Andrea S, Segatto I, Canzonieri V, et al. Genetic characterization of p27^kip1^ and stathmin in controlling cell proliferation in vivo. Cell Cycle. 2014;13(19):3100–11. doi: 10.4161/15384101.2014.949512. [PMID: 25486569].

34. Fabris L, Berton S, Pellizzari I, Segatto I, D’Andrea S, Armenia J, et al. p27^kip1^ controls H-Ras/MAPK activation and cell cycle entry via modulation of MT stability. Proc Natl Acad Sci USA. 2015 Nov 10;112(45):13916–21. doi: 10.1073/pnas.1508514112. [PMID: 26512117].

35. Vlach J, Hennecke S, Amati B. Phosphorylation-dependent degradation of the cyclin-dependent kinase inhibitor p27. EMBO J. 1997 Sep 1;16(17):5334–44. doi: 10.1093/emboj/16.17.5334. [PMID: 9311993].

36. Zong H, Espinosa JS, Su HH, Muzumdar MD, Luo L. Mosaic analysis with double markers in mice. Cell. 2005 May 6;121(3):479–92. doi: 10.1016/j.cell.2005.02.012. [PMID: 15882628].

37. Tang SH, Silva FJ, Tsark WM, Mann JR. A Cre/loxP-deleter transgenic line in mouse strain 129S1/SvimJ. Genesis. 2002 Mar;32(3):199–202. [PMID: 11892008].

38. Espinosa JS, Tea JS, Luo L. Mosaic analysis with double markers (MADM) in mice. Cold Spring Harb Protoc. 2014 Feb 1;2014(2):182–9. doi: 10.1101/pdb.prot080366. [PMID: 24492775].

39. Zong H. Generation and applications of MADM-based mouse genetic mosaic system. Methods Mol Biol. 2014; 1194:187–201. doi: 10.1007/978-1-4939-1215_10. [PMID: 25064104].

40. Muzumdar MD, Luo L, Zong H. Modeling sporadic loss of heterozygosity in mice by using mosaic analysis with double markers (MADM). Proc Natl Acad Sci U S A. 2007 Mar 13;104(11)al:4495-500. doi: 10.1073/pnas.0606491104. [PMID: 17360552].

41. Lorincz A, Nusser Z. Molecular identity of dendritic voltage-gated sodium channels. Science. 2010 May 14;328:906–9. doi: 10.1126/science.1187958. [PMID: 20466935].

42. Espinosa JS, Luo L. Timing neurogenesis and differentiation: Insights from quantitative clonal analyses of cerebellar granule cells. J Neurosci. 2008 Mar 5;28(10):2301–12. doi: 10.1523/JNEUROSCI.5157-07.2008. [PMID: 18322077].

43. Schindelin J, Arganda-Carreras I, Frise E, Kaynig V, Longair M, Pietzsch T, et al. Fiji: an open-source platform for biological-image analysis. Nat Methods. 2012 June 28;9(7):676–682. doi: 10.1038/nmeth.2019. [PMID:22743772].

44. Ester M, Kriegel H-P, Sander J, Xu X. A density-based algorithm for discovering clusters in large spatial databases with noise. In: Simoudis E, Han J, Fayyad UM, editors. Proceedings of the Second International Conference on Knowledge Discovery and Data Mining (KDD-96); AAAI Press; 1996. p. 226–231. CiteSeerX 10.1.1.121.9220. [ISBN 1-57735-004-9].

45. Germani F, Bergantinos C, Johnson LA. Mosaic analysis in *Drosophila*. Genetics. 2018 Feb;208(2):473–490. doi:10.1534/genetics.117.300256. Review. [PMID:29378809].

46. Baldassarre G, Belletti B, Nicoloso MS, Schiappacassi M, Vecchione A, Spessotto P, et al. p27^Kip1^-stathmin interaction influences sarcoma cell migration and invasion. Cancer Cell. 2005 Jan;7(1):51–63. doi: 10.1016/j.ccr.2004.11.025. [PMID: 15652749].

47. Besson A, Gurian-West M, Schnidt A, Hall A, Roberts JM. p27Kip1 modulates cell migration through the regulation of RhoA activation. 2004 Apr 15;18(8):862–76. doi: 10.1101/gad.1185504. [PMID: 15078817].

48. Olson MF, Ashworth A, Hall A. An essential role for Rho, Rac, and Cdc42 GTPases in cell cycle progression through G1. Science. 1995 Sep 1;269(5228):1270–2. [PMID: 7652575]

49. Yamamoto M, Marui N, Sakai T, Morii N, Kozaki S, Ikai K, Imamura S, Narumiya S. ADP-ribosylation of the rhoA gene product by botulinum C3 exoenzyme causes Swiss 3T3 cells to accumulate in the G1 phase of the cell cycle. Oncogene. 1993 Jun;8(6):1449–55. [PMID: 8502473].

50. Coleman ML, Marshall CJ, Olson MF. RAS and RHO GTPases in G1-phase cell-cycle regulation. Nat Rev Mol Cell Biol. 2004 May;5(5):355–66. [PMID: 15122349].

51. Croft DR, Olson MF. The Rho GTPase effector ROCK regulates cyclin A, cyclin D1, and p27Kip1 levels by distinct mechanisms. Mol Cell Biol. 2006 Jun;26(12):4612–27. [PMID: 16738326].

52. Okumura N, Ueno M, Koizumi N, Sakamoto Y, Hirata K, Hamuro J, Kinoshita S. Enhancement on primate corneal endothelial cell survival in vitro by a ROCK inhibitor. Invest Ophthalmol Vis Sci. 2009 Aug;50(8):3680–7. doi: 10.1167/iovs.08-2634. [PMID: 19387080].

53. Okumura N, Nakano S, Kay EP, Numata R, Ota A, Sowa Y, Sakai T, Ueno M, Kinoshita S, Koizumi N. Involvement of cyclin D and p27 in cell proliferation mediated by ROCK inhibitors Y-27632 and Y-39983 during corneal endothelium wound healing. Invest Ophthalmol Vis Sci. 2014 Jan 15;55(1):318–29. doi: 10.1167/iovs.13-12225. [PMID: 24106120].

54. Okumura N, Koizumi N, Kay EP, Ueno M, Sakamoto Y, Nakamura S, Hamuro J, Kinoshita S. The ROCK inhibitor eye drop accelerates corneal endothelium wound healing. Invest Ophthalmol Vis Sci. 2013 Apr 3;54(4):2493–502. doi: 10.1167/iovs.12-11320. [PMID: 23462749].

55. Okumura N, Inoue R, Okazaki Y, Nakano S, Nakagawa H, Kinoshita S, Koizumi N. Effect of the Rho kinase inhibitor Y-27632 on corneal endothelial wound healing. Invest Ophthalmol Vis Sci. 2015 Sep;56(10):6067–74. doi: 10.1167/iovs.15-17595. [PMID: 26393474].

56. Okumura N, Okazaki Y, Inoue R, Kakutani K, Nakano S, Kinoshita S, Koizumi N. Effect of the Rho-associated kinase inhibitor eye drop (Ripasudil) on corneal endothelial wound healing. Invest Ophthalmol Vis Sci. 2016 Mar;57(3):1284–92. doi: 10.1167/iovs.15-18586. [PMID: 26998714].

57. Meekins LC, Rosado-Adames N, Maddala R, Zhao JJ, Rao PV, Afshari NA. Corneal endothelial cell migration and proliferation enhanced by Rho kinase (ROCK) inhibitors in in vitro and in vivo models. Invest Ophthalmol Vis Sci. 2016 Dec 1;57(15):6731–38. doi: 10.1167/iovs.16-20414. [PMID: 27951595].

58. Zhu YT, Chen HC, Chen SY, Tseng SC. Nuclear p120 catenin unlocks mitotic block of contact-inhibited human corneal endothelial monolayers without disrupting adherent junctions. J Cell Sci. 2012 Aug 1;125(Pt 15):3636–48. doi: 10.1242/jcs.103267. [PMID: 22505615].

59. McAllister SS, Becker-Hapak M, Pintucci G, Pagano M, Dowdy SF. Novel p27^kip1^ C-terminal scatter domain mediates Rac-dependent cell migration independent of cell cycle arrest functions. Mol Cell Biol. 2003 Jan;23(1):216–28. [PMID: 12482975].

60. Nagahara H, Vocero-Akbani AM, Snyder EL, Ho A, Latham DG, Lissy NA, Becker-Hapak M, Ezhevsky SA, Dowdy SF. Transduction of full=length TAT fusion proteins into mammalian cells: TAT-p27Kip1 induces cell migration. Nat Med. 1998 Dec;4(12):1449–52. doi: 10.1038/4042. [PMID: 9846587].

61. Kawauchi T, Shikanai M, Kosodo Y. Extra-cell cycle regulatory functions of cyclin-dependent kinases (CDK) and CDK inhibitor proteins contribute to brain development and neurological disorders. Genes Cells. 2013 Mar;18(3):176–94. doi: 10.1111/gtc.12029. doi: 10.1111/gtc.12029. [PMID: 23294285].

62. Nguyen L, Besson A, Heng JI, Schuurmans C, Teboul L, Parras C, Philpott A, Roberts JM, Guillemot F. p27kip1 independently promotes neuronal differentiation and migration in the cerebral cortex. Genes Dev. 2006 Jun 1;20(11):1511–24. doi: 10.1101/gad.377106. [PMID: 16705040].

63. Sheaff RJ, Groudine M, Gordon M, Roberts JM, Clurman BE. CyclinE-CDK2 is a regulator of p27Kip1. Genes Dev. 1997 Jun 1;11(11):1464–78. [PMID: 9192873].

