## Supplemental Figures 1 and 2 for "A non-canonical role for p27^Kip1^ in restricting proliferation of corneal endothelial cells during development"

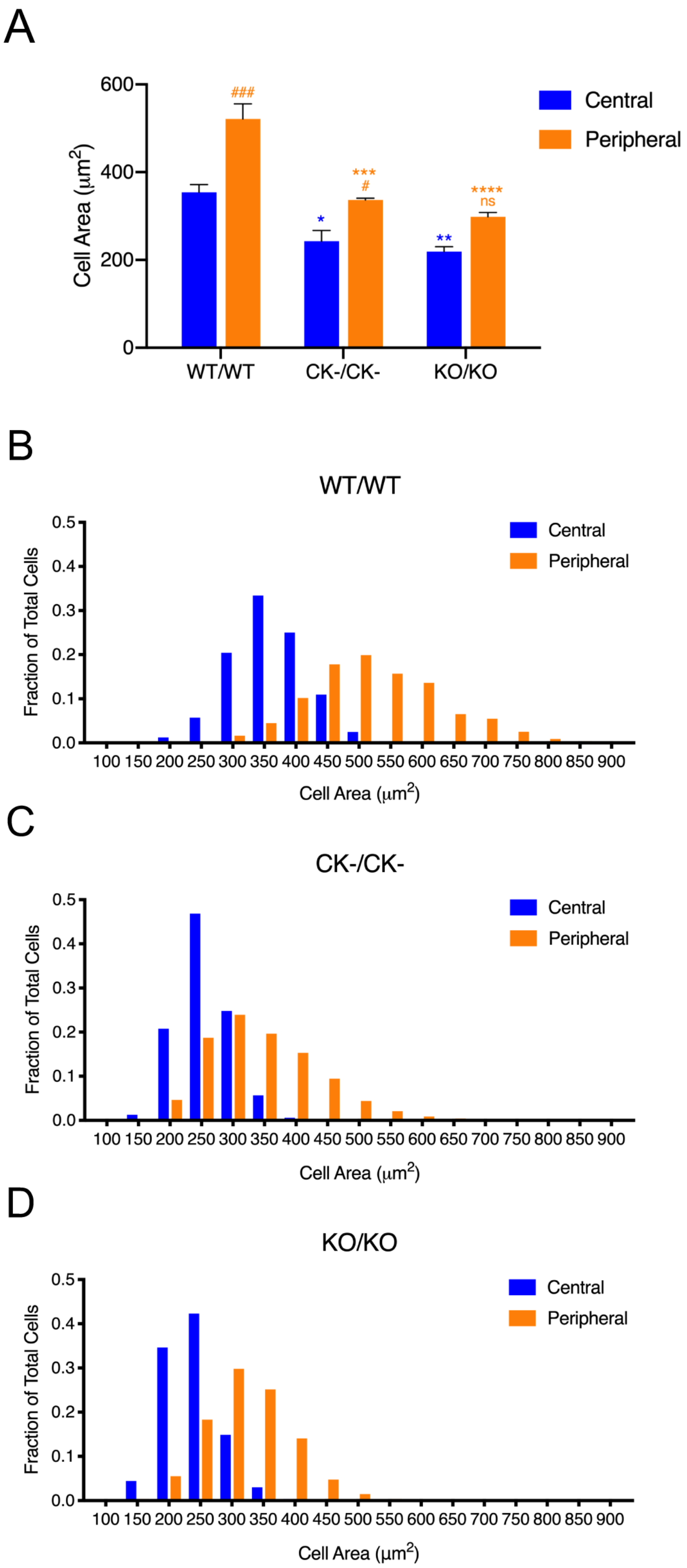

A
600
400
200
0
Cell Area (μm2)
###
***
#
****
ns
*
**
WT/WT
CK-/CK-
KO/KO
Central
Peripheral
B
WT/WT
Fraction of Total Cells
0.5
0.4
0.3
0.2
0.1
0.0
100 150 200 250 300 350 400 450 500 550 600 650 700 750 800 850 900
Cell Area (μm2)
Central
Peripheral
C
CK-/CK-
Fraction of Total Cells
0.5
0.4
0.3
0.2
0.1
0.0
100 150 200 250 300 350 400 450 500 550 600 650 700 750 800 850 900
Cell Area (μm2)
Central
Peripheral
D
KO/KO
Fraction of Total Cells
0.5
0.4
0.3
0.2
0.1
0.0
100 150 200 250 300 350 400 450 500 550 600 650 700 750 800 850 900
Cell Area (μm2)
Central
Peripheral

Supplement Figure 1. Measurement of apical cell areas in wild-type (WT/WT) and mutant (CK−/CK− and KO/KO) endothelia. (A) Averaged data. Declines in individual cell area are evident for central and peripheral regions of both mutants, relative to the wild-type control. (B-D) Histograms plotting cell area distributions. In *p27*$^{+/+}$ monolayers, the size ranges of regional populations are relatively broad and partially overlapping. Expression of mutant alleles leads to shifts in cell sizes to smaller values in both central and peripheral regions, as well as more compressed distributions and greater overlap. Data in (A) represent means ± SEM (n = 5). Ordinary two-way ANOVA followed by Tukey's HSD test was performed. * and ** indicate $p<0.05$ and $p<0.005$ by comparison to wild-type (central), while *** and **** indicate $p<0.0005$ and $p<0.0001$ by comparison to wild-type (peripheral). # and ### indicate $p<0.05$ and $p<0.001$ by comparison to central regions. ns indicates not significant.

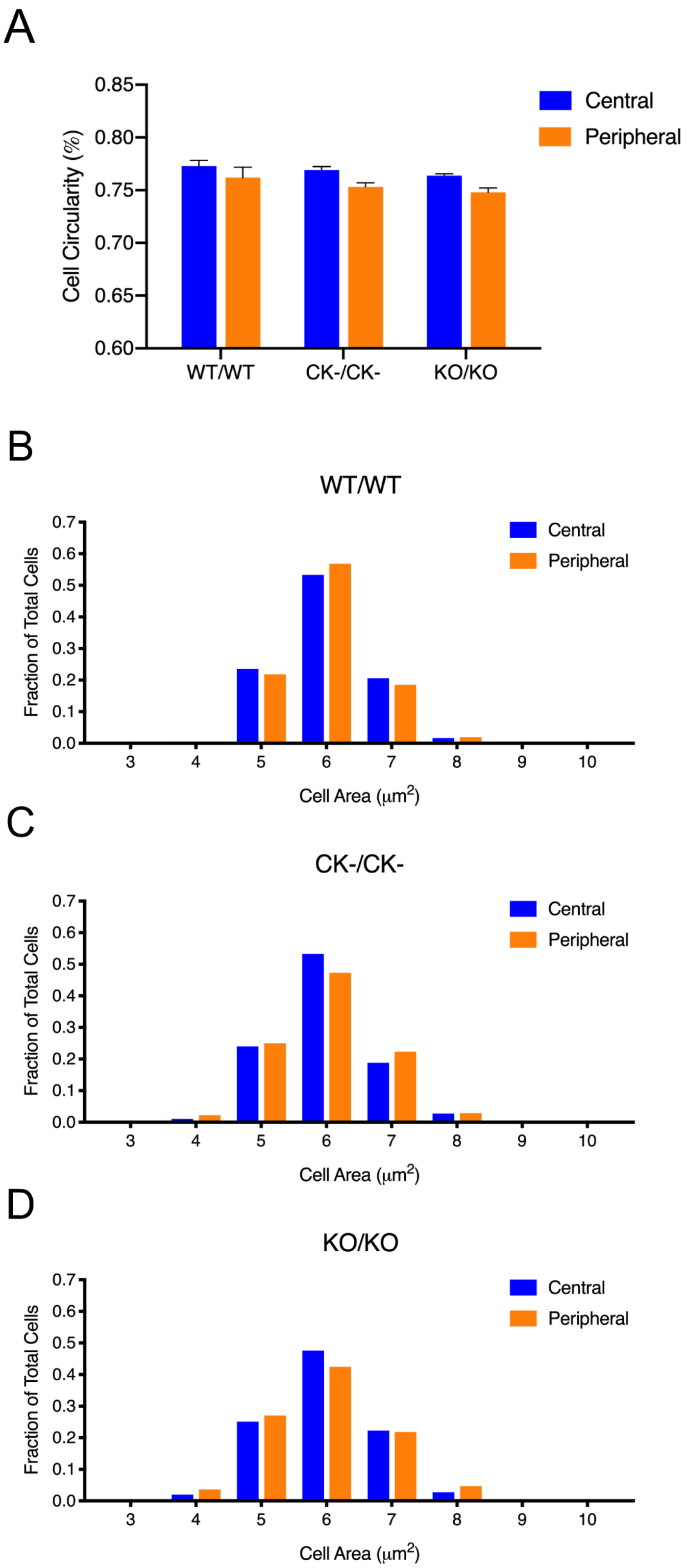
A
Cell Circularity (%)
0.85
0.80
0.75
0.70
0.65
0.60
WT/WT
CK-/CK-
KO/KO
Central
Peripheral
B
WT/WT
Fraction of Total Cells
Cell Area (μm2)
C
CK-/CK-
D
KO/KO

Supplement Figure 2. Shape and neighbor analysis of wild-type (WT/WT) and mutant (CK−/CK− and KO/KO) endothelial cells. (A) Averaged circularity data. Peripheral cells exhibit a small, but significant, decline in circularity across all genotypes ($p<0.01$). However, no difference is observed in either of the two regional cell populations when comparing wild-type and mutant monolayers. (B-D) Histogram plots of nearest neighbor distributions. Quantitatively similar numbers of neighbors are seen for all genotypes.
